# A novel antigenic site spanning domains I and III of the Zika virus envelope glycoprotein is the target of strongly neutralizing human monoclonal antibodies

**DOI:** 10.1101/2020.12.21.423896

**Authors:** Stephen Graham, Huy A. Tu, Benjamin D. McElvany, Nancy R. Graham, Ariadna Grinyo, Edgar Davidson, Benjamin J. Doranz, Sean A. Diehl, Aravinda M. de Silva, Alena Janda Markmann

## Abstract

Zika virus (ZIKV), a mosquito-transmitted flavivirus, caused a large epidemic in Latin America between 2015 and 2017. Effective ZIKV vaccines and treatments are urgently needed to prevent future epidemics and severe disease sequelae. People infected with ZIKV develop strongly neutralizing antibodies linked to viral clearance and durable protective immunity. To understand mechanisms of protective immunity and to support the development of ZIKV vaccines, here we characterize the properties of a strongly neutralizing antibody, B11F, isolated from a recovered ZIKV patient. Our results indicate that B11F targets a complex epitope on the virus that spans domains I and III of the envelope glycoprotein. While previous studies point to quaternary epitopes centered on domain II of ZIKV E glycoprotein as targets of strongly neutralizing and protective human antibodies, we uncover a new site spanning domain I and III as a target of strongly neutralizing human antibodies.

**Importance:** People infected with Zika virus develop durable neutralizing antibodies that prevent repeat infections. In the current study, we characterize a ZIKV-neutralizing human monoclonal antibody isolated from a patient after recovery. Our studies establish a novel site on the viral envelope targeted by human neutralizing antibodies. Our results are relevant to understanding how antibodies block infection and for guiding the design and evaluation of candidate vaccines.

## Introduction

ZIKV is a mosquito-borne flavivirus responsible for recent large epidemics accompanied by severe clinical manifestations such as Guillain-Barre syndrome and congenital birth defects (1). The ZIKV epidemic in South America in 2015 highlighted the need to understand immune mechanisms of protective immunity to guide the development of vaccines and other counter measures (2). Among flaviviruses, ZIKV is most closely related to the four dengue viruses (DENV 1-4). Recent setbacks faced by DENV vaccine developers highlight the importance of a deeper understanding of immune-protective antigenic targets to pathogenic flaviviruses (3, 4).

In humans, infection with a single flavivirus is known to induce long-term, likely life-long, adaptive immune protection. An important component of the long-term protective immune response to flaviviral infections is the production of potently neutralizing antibodies (5). These durable protective antibodies generated after flaviviral infection, are continuously secreted by long-lived plasma cells in the bone marrow and produced during antigenic recall by memory B cells (MBC) residing in lymphoid organs. Identifying the viral binding sites, or antigenic targets of MBC-derived neutralizing antibodies helps us understand how the immune response prevents repeated viral infections by the same virus (6). Immunogenic epitopes can be used to directly inform vaccine and also aid in diagnostic design (7, 8).

The main target of human antibodies that neutralize flaviviruses is the envelope (E) glycoprotein that covers the surface of the virion. Each E glycoprotein monomer contains three domains: domain I (EDI), II (EDII), and III (EDIII), and a fusion loop at the tip of domain II (Figure 4) (9). E glycoproteins form stable homodimers and 90 dimers assemble to form the outer envelope of the infectious virus. Primary flavivirus infections stimulate cross-reactive (CR) antibodies that target epitopes conserved between closely related flaviviruses as well as type-specific (TS) antibodies that bind to unique epitopes on the infecting virus. CR antibodies do not reliably confer durable cross-protective immunity after a primary infection, most likely because they bind with low affinity to conserved epitopes that are not well exposed on the viral surface (10). In contrast, TS antibodies are often strongly neutralizing and linked to long-term protection from re-infection by the same flavivirus (2).

Several groups have isolated a few strongly neutralizing human monoclonal antibodies (mAbs) from MBCs. The most potent antibodies have been mapped to complex epitopes centered on domain II of the E glycoprotein with footprints that span two or more E glycoproteins (11-13). Recent studies have also identified a few strongly neutralizing ZIKV mAbs that target epitopes in EDIII (14-17), although this domain does not appear to be a major target of serum polyclonal neutralizing antibody responses (7). Using mAbs isolated from individuals who have recovered from Zika, here we define a novel antigenic site, between domains I and III of the ZIKV E glycoprotein, targeted by human antibodies that strongly neutralize ZIKV.

## Results

### Isolation of a ZIKV type-specific and strongly neutralizing mAb B11F

Subject DT172 was a US traveler who acquired a self-limited and uncomplicated ZIKV infection while traveling through Nicaragua and Colombia in 2015, early in the South American ZIKV epidemic. The subject had no neutralizing antibody titers to DENV 1-4 (< 20), and a neutralization titer of 1:794 to ZIKV. As previously reported (11), when MBCs collected 3 months after recovery were immortalized by the 6XL method and tested, 0.9% of the cells were observed to be producing ZIKV-binding antibodies. To isolate ZIKV neutralizing mAbs, we single-cell sorted MBCs and tested individual clones for ZIKV binding and neutralizing antibody as previously described(11). We identified and sequenced a single IgG1 clone, named B11F (Table 1), that strongly neutralized ZIKV (FRNT_50_ = 3.22 ng/mL) (Figure 1A).

**Table 1.** Identification of residues critical for ZIKV mAb B11F binding. Binding data for B11F and A9E at all ZIKV E protein clones identified as critical for B11F binding. mAb reactivity for each mutant are expressed as percent of binding to wild type ZIKV prME, with ranges (half of the maximum minus minimum values) in parentheses. At least two replicate values were obtained for each experiment.

| Mutation | B11F Binding (%WT) | A9E Binding (%WT) |
| --- | --- | --- |
| R138A | -1.4 (5.7) | 78.2 (1.1) |
| K166A | 2.4 (1.4) | 93.4 (33.7) |
| M140A | 3.1 (2.9) | 107.6 (12.5) |
| R164A | 12.7 (7.0) | 105.2 (14.5) |
| D161A | 13.3 (6.5) | 40.5 (10.1) |
| K281A | 13.5 (3.4) | 77.1 (3.6) |
| T156A | 23.4 (2.9) | 59.7 (16.8) |

**Figure 1.**
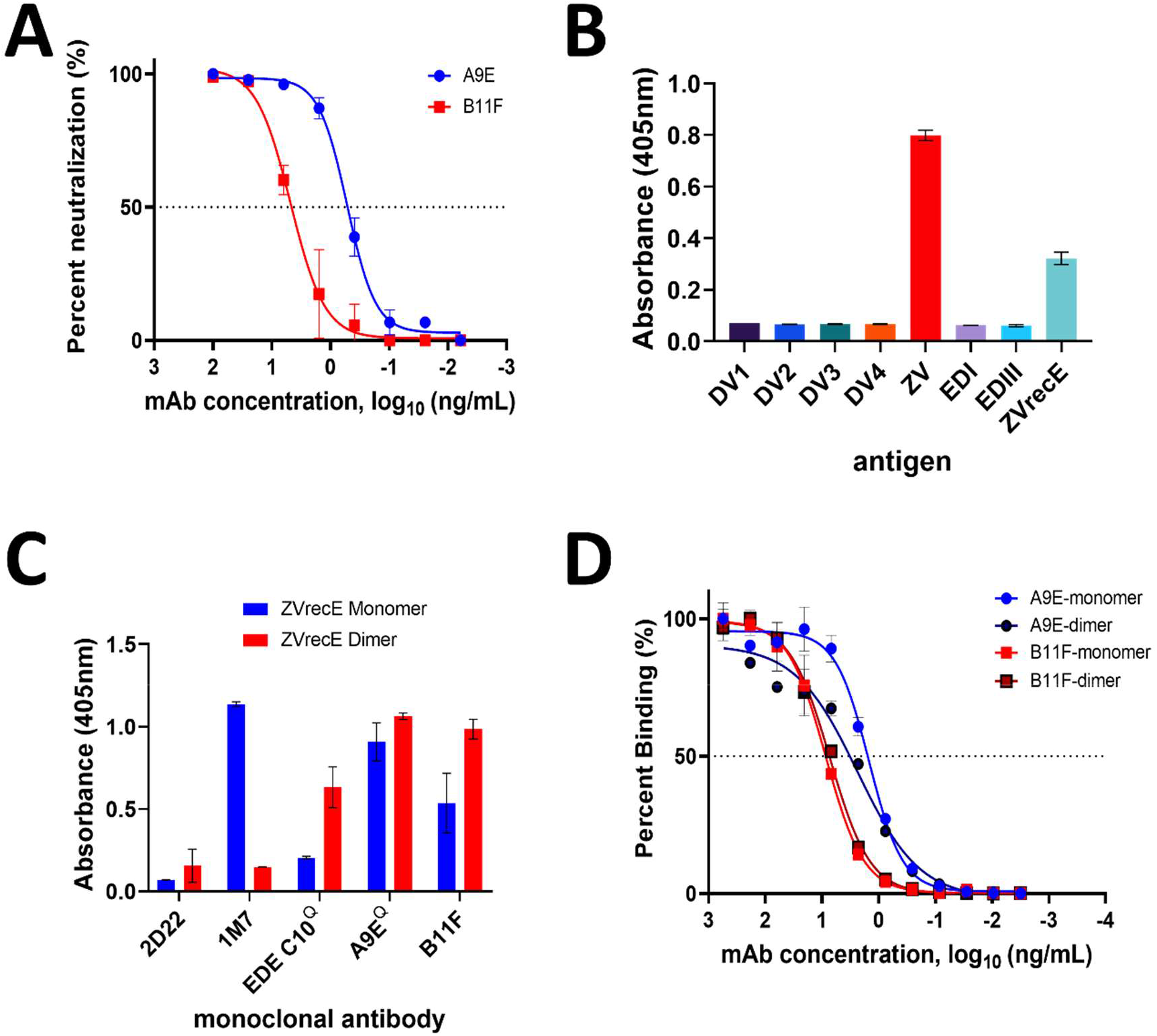
B11F ELISA binding specificity and virus neutralization. A) ZIKV neutralization by B11F and A9E. Each value is the average of duplicate wells. 50% neutralization occurred at concentrations of 3.22 ng/mL for B11F (open squares) and 0.33 ng/mL for A9E (open circles). Graph is a representative of three independent experiments. B) B11F mAb binding ELISA using whole virions, recombinant ZIKV E protein and domains EDI and EDIII. C) B11F binding to ZIKV E protein monomer and dimer by capture ELISA. 2D22 is a DENV-2 specific mAb, 1M7 is a fusion-loop binding pan-flaviviral antibody, and EDE C10 binds a quaternary epitope only present on dimer antigen. For B and C, each value represents an average of duplicate wells, background absorbance is 0.1 optical density (OD) units, and graph representative of at least two independent experiments. D) B11F and A9E binding to monomer and dimer forms of ZIKV E protein. A9E+monomer (EC50 = 1.7ng/mL; blue circle), A9E+dimer (EC50 = 2.1ng/mL; black circle), B11F+monomer (EC50 = 7.6ng/mL; red square), B11F+dimer (EC50 = 5.8ng/mL; black square). EC50 values are an average of two independent experiments. Each value is the average of duplicate wells.

Recombinant produced B11F IgG1 bound to ZIKV but not DENV1-4 (Figure 1B). The antibody also bound to the full length ectodomain (ZVrecE, wild-type protein) but not domains I or III of ZIKV E glycoprotein (Figure 1B). In solution at 37°C, the ZVrecE glycoprotein is in an equilibrium that greatly favors monomers over homodimers (9). We compared the binding of several ZIKV-specific human mAbs, including B11F to ZVrecE monomers and stable homodimers (stabilized by the introduction of an intermolecular disulfide bond (18). Control antibodies that preferentially bound monomers (pan-flaviviral fusion-loop targeting mAb 1M7) or homodimers (quaternary epitope-specific mAb EDE C10) confirmed the oligomeric state of our antigens (Figure 1C). B11F bound similarly to ZVrecE monomers and homodimers (Figure 1D). We conclude that B11F is a potently neutralizing antibody that binds to a ZIKV type-specific epitope that is displayed on the ectodomain of ZIKV E glycoprotein but not on domains I and III alone. This is similar to another human mAb designated A9E that we recently isolated from a traveler who was infected with ZIKV in Brazil in 2017 (Figure 1) (11).

### Blockade of binding (BOB) assays with B11F and other well-characterized ZIKV-specific human mAbs

To further explore how the ZIKV E glycoprotein binding site of B11F is related to known antibody epitopes on ZIKV, we performed antibody competition assays (BOB assays) with B11F versus fourteen ZIKV-specific mAbs with mapped epitopes. We compared the ability of each ZIKV-specific mAb to block the binding of labelled B11F to intact ZIKV virions captured on an ELISA plate (Figure 2). Out of the fourteen antibodies tested, only the A9E mAb blocked the binding of B11F to ZIKV. While the full epitope of A9E has not been mapped yet, our previous studies demonstrate that A9E targets an epitope centered on EDI that extends towards EDIII on a single E glycoprotein monomer (11). This supports the evidence that the B11F mAb targets the EDI-EDIII domain on the ZIKV E glycoprotein.

**Figure 2.**
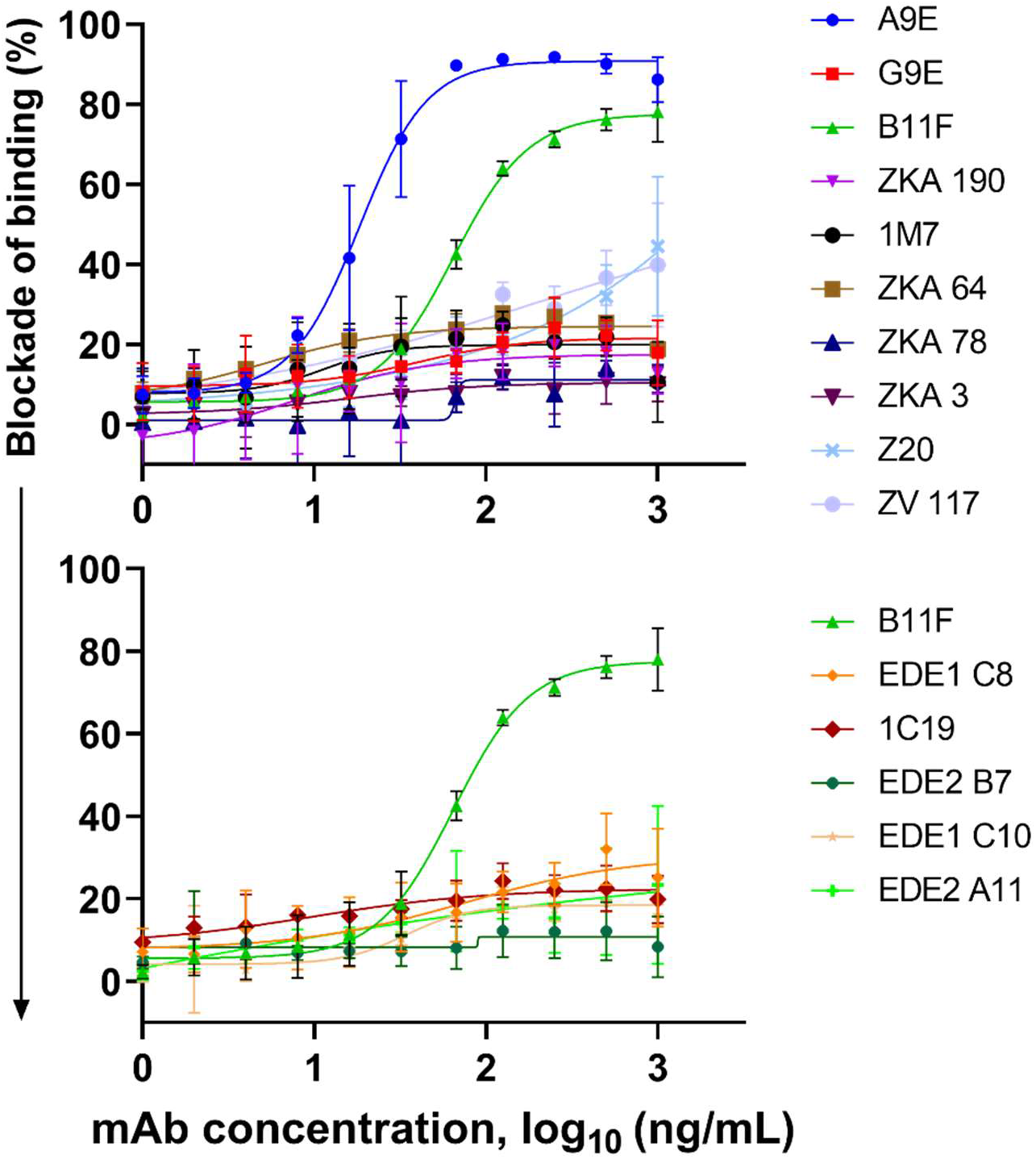
Zika virus blockade of binding ELISA results. Top panel) B11F blockade with Zika-specific monoclonal antibodies. Bottom panel) B11F blockade with dengue virus-specific monoclonal antibodies. Here B11F is held at constant concentration, the x-axis shows the varying concentration of the competing monoclonal antibody. Error bars represent averaged data sets from two independent experiments.

### Identification of ZIKV neutralization escape mutants

The epitopes of mAbs that neutralize flaviviruses can be mapped by passaging virus in the presence of the mAb under study to select for mutations that prevent antibody binding and neutralization. We previously reported on specific mutations on EDI (G182D) and EDIII (V364I) that led to neutralization escape from mAb A9E (11) (Figure 4). Mutations that led to binding and neutralization escape from mAb A9E moderately reduced binding as well as neutralization potency of B11F (B11F FRNT_50_ = ∼3 ng/mL (wild-type) → ∼34 ng/mL (escape mutant)) (Figure 3A and B). We also passaged ZIKV in the presence of mAb B11F and isolated an escape mutant virus that was able to replicate in the presence of B11F. The B11F escape mutant virus has a mutation at position M345I, which is buried in the core of EDIII (Figure 4A). The M345I mutation, which prevented binding and neutralization by mAb B11F, had a moderate effect on mAb A9E binding, and a 10-fold reduction in its neutralization potency when compared to wild-type virus neutralization (A9E FRNT_50_ = ∼0.3 ng/mL (wild-type) → ∼6 ng/mL (escape mutant)) (Figures 3C and 3D). These results are consistent with human mAbs B11F and A9E having over-lapping but distinct epitopes.

**Figure 3.**
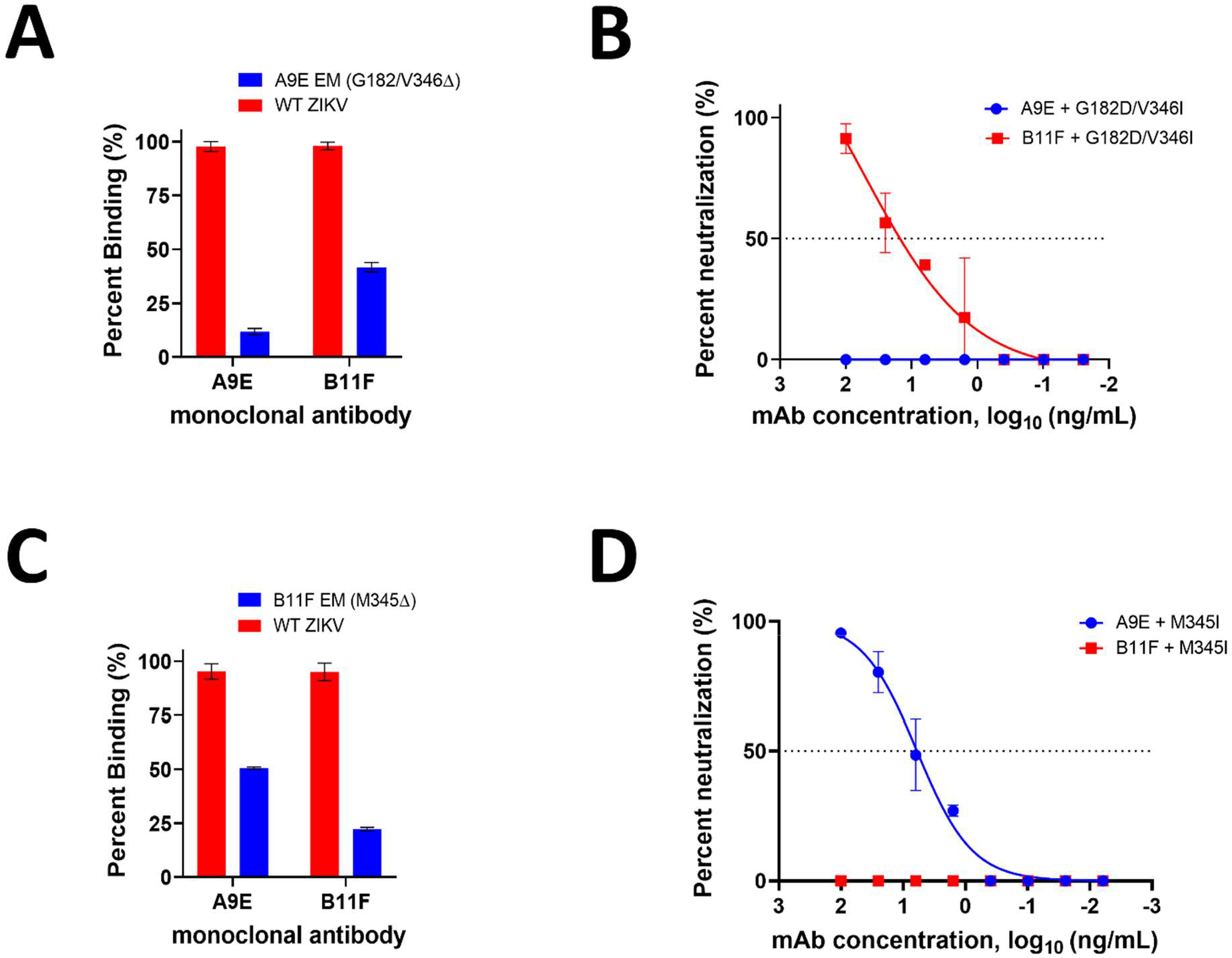
Binding and neutralization of escape mutant viruses. A) Whole virion capture ELISA binding results for the A9E escape mutant virus. Percent binding values represent the average of duplicate wells. B) A9E escape mutant virus neutralization assay, WT = wild-type ZIKV. Mutations shown are those located on the A9E escape mutant virus. C) Whole virion capture ELISA binding results for the B11F escape mutant. D) B11F escape mutant virus neutralization assay. Mutations shown are those located on the B11F escape mutant virus. All graphs shown are representative of at least two independent experiments.

**Figure 4.**
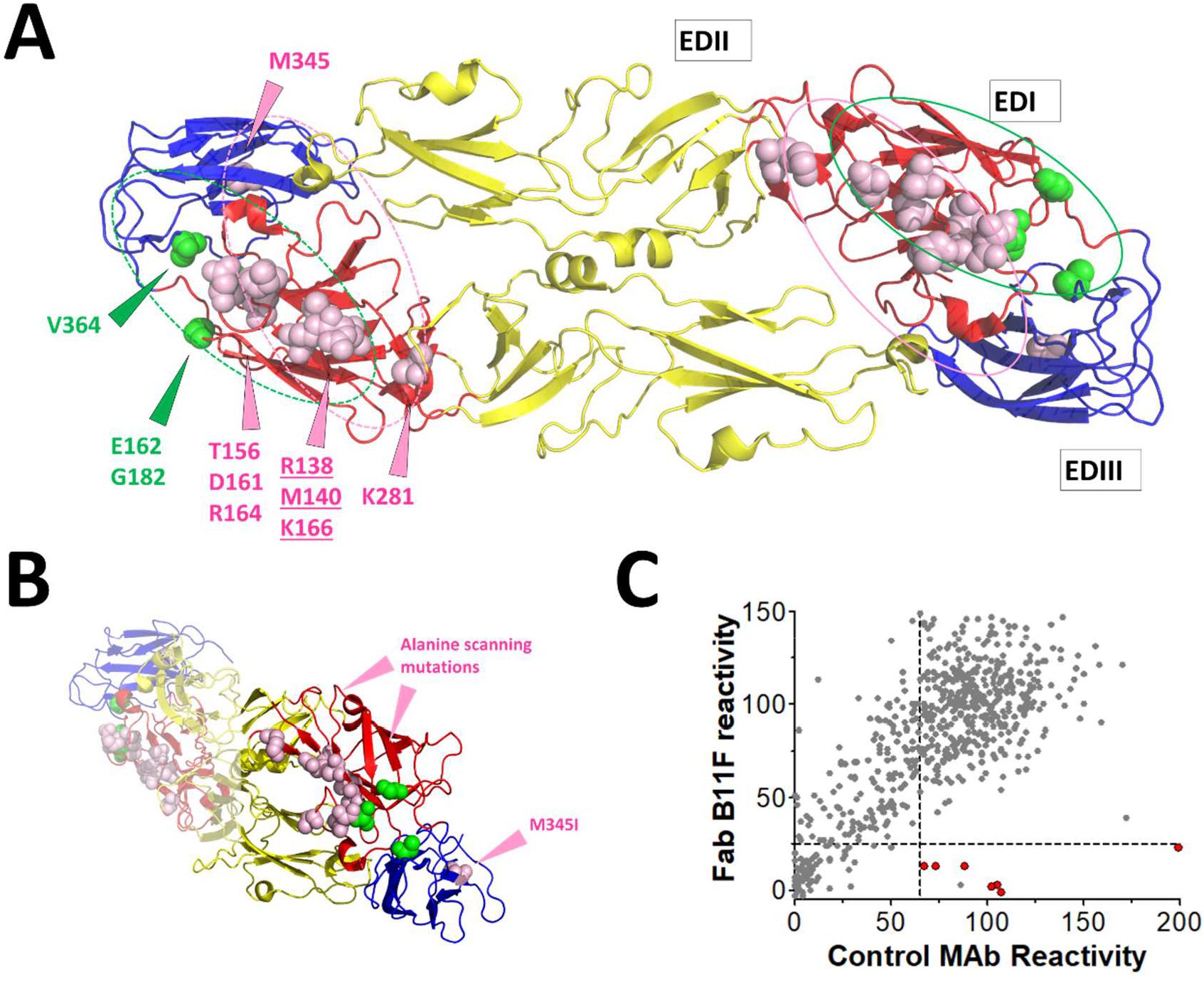
Epitope mapping analysis of B11F and A9E antibodies. A) Zika virus envelope protein dimer (PDB: 5IRE) with domains labeled and color-coated. Location of B11F viral escape mutation (M345; pink spheres within EDIII), and alanine scanning mutations (pink spheres on EDI; underlined residues have largest contribution to binding). Locations of A9E escape mutations and alanine scanning mutations (green spheres). Putative B11F mAb footprint in pink, putative A9E mAb footprint in green. B) Side/edge view displaying distance between B11F escape mutation M345I and the B11F mutations identified by alanine scanning. Distance between the closest atoms of M345 and the alanine scanning mutant residues for B11F are M345-N and D161-O is approximately 27 Angstroms. C) Critical amino acid residues for B11F Fab binding to ZIKV envelope glycoprotein as determined by alanine scanning shotgun mutagenesis. This plot shows B11F Fab binding to the mutants versus a set of control monoclonal antibodies. Red circles correspond to alanine mutants that reduce B11F Fab binding compared to control monoclonal antibodies.

### Alanine Scanning Mutagenesis for epitope mapping

We used a ZIKV prM/E glycoprotein expression library with single alanine mutations to identify mutations that reduced or eliminated mAb B11F Fab binding. This library and approach has been extensively validated for mapping ZIKV binding antibodies (11, 13). Mutations at E glycoprotein residues Arg138, Thr156, Met140, Asp161, Arg164, Lys166 and Lys281 selectively reduced B11F binding, while retaining the overall structural integrity of the glycoprotein (Figure 4A). Mutations at residues M140, K166, and R138 had the largest impact on B11F binding, and very little to no impact on the A9E mAb (Table 1). All the residues identified by alanine scanning mutagenesis are surface exposed on EDI. The shortest distance between the escape virus mutation M345I and an alanine scan-identified mutation was between M345-N and D161-O, which is a large distance of approximately 27 Angstroms (Figure 4B).

### Antibody Genetics of ZIKV type-specific and potently neutralizing human mAbs

The human mAb A9E is a ZIKV type-specific and strongly neutralizing antibody that binds to a distinct epitope centered on E glycoprotein domain I-III. The mAb B11F was isolated from a different individual and has an epitope that partially overlaps with the A9E epitope. Both of these mAbs are IgG1 antibodies with a lambda light chain (Table 2). Compared to eight other MBC-derived monoclonal antibodies that neutralize Zika virus and have known V-D-J gene usage, B11F is the only one using the V5-10 heavy chain gene locus (11, 17, 19-24). Heavy chain V-gene usage is also different between B11F and A9E (Table 2). The B11F light chain gene locus on the other hand, V2-14*01 is also used by at least three other Zika neutralizing antibodies: A9E, G9E and C10 (11, 23). All four of these monoclonal antibodies share a similar CDRL3 sequence. Notably, B11F has fewer non-silent somatic hypermutations than A9E in both V and H-genes, which suggests a higher degree of somatic hypermutation and may explain the stronger neutralization potency of A9E compared to that of B11F despite a similar E glycoprotein epitope (Table 2). Stronger neutralization potency may suggest that the A9E mAb has higher affinity for ZIKV E glycoprotein dimers than the B11F mAb, on a single molecule level, and we plan to test this hypothesis in the future. Despite their different origins, mAbs B11F and A9E have similar CDRH3 and CDRL3 sequences, which is consistent with their binding to a shared epitope on the E glycoprotein.

**Table 2.**
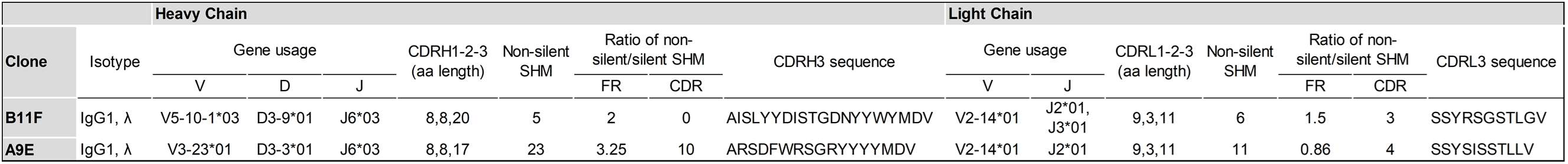
Comparison of sequence and IgG characteristics of Zika monoclonal antibodies A9E and B11F. (From https://www.ncbi.nlm.nih.gov/igblast/). (A9E data from Collins *et al*.(11))

## Discussion

In this study, we have identified a new region on the ZIKV envelope glycoprotein targeted by two strongly neutralizing human MBC-derived monoclonal antibodies that have been isolated from two separate individuals. Both mAbs bound to the monomeric form of the E glycoprotein indicating that most of the footprint is contained within a single E protein molecule. In this regard, both antibodies are different from other human mAbs that strongly neutralize flaviviruses and bind to quaternary epitopes that span two more E molecules on the viral surface (11, 12, 19).

By passaging ZIKV in the presence of mAb B11F to select for escape mutations, we identified a residue buried within the core of EDIII (M345) that was critical for binding and neutralization. We hypothesize that this mutation results in allosteric changes within EDIII that influence the surface epitope on the EDIII or EDI domains that interacts with B11F during virus binding and neutralization. The A9E monoclonal antibody showed decreased binding and neutralization, though still potent, to the B11F escape mutant. Similarly, mutations in the EDI/III hinge region that promoted neutralization escape from A9E had a significant impact on B11F binding as well as neutralization. These observations indicate that though the footprints of both of these monoclonal antibodies are similar and likely overlap, they are not identical. Binding and viral escape mutation studies, though good surrogates for predicting viral epitopes of potent mAbs, are a limitation here in that they do not give us direct structural or functional data, which will have to be done in future studies to fully understand the epitopes and mechanisms of these mAbs.

While B11F did not bind to EDI alone, we predict the footprint of this antibody to be mainly contained on EDI because of the many mutations in this domain identified by alanine scanning mutagenesis (Figure 4). Similarly, the majority of the epitope for A9E appears to be centered on EDI because the antibody was able to bind to EDI alone produced as a recombinant antigen. While viral mutations in EDIII (B11F) or the linker region between EDI and III (A9E) resulted in complete neutralization escape, neither mAb bound to EDIII when separated from the rest of the E glycoprotein. This indicates that both antibodies require interaction with EDI and an adjacent EDIII domain for functional neutralization of ZIKV. However, the expanded footprints of the two antibodies differ because EDIII mutations had distinct phenotypes for each mAb. Residues identified by alanine-scanning as important are thought to be those most energetically important for antibody binding (31, 32). Although we identified residues for B11F only in DI, it is possible that DIII has epitope contact residues that not energetically important for binding, but subject to perturbation by escape mutation. Furthermore, the three alanine scanning mutations with the highest energetic importance for B11F binding, had little effect on A9E binding. Taken together, the mutagenesis and escape mutant studies reveal that B11F and A9E rely on different points of contact on the ZIKV E glycoprotein surface, with A9E covering the outer portion of the EDI-EDIII hinge, and B11F shifted inward covering EDI, with possible contacts on EDIII and EDII as well.

Previous studies from our group and other groups indicate that quaternary structure epitopes centered on ZIKV EDII with footprints that expand into adjacent molecules of E glycoprotein homodimers and higher order structures act as targets of strongly neutralizing and protective human antibodies (11, 12, 19). In this study we propose that the A9E and B11F monoclonal antibodies define a new antigenic region spanning EDI and EDIII within a single E protein targeted by the neutralizing human antibody response to ZIKV. The EDI-EDIII interface is an important immunogenic epitope on the ZIKV E glycoprotein that can be leveraged for ZIKV vaccine design. Furthermore, both A9E and B11F have the potential to be used as future therapeutic antibodies for treatment of ZIKV or as prophylaxis during a ZIKV outbreak.

## Materials and Methods

### Human subjects and biospecimen collection

Whole blood donations were obtained from fully consented volunteer travelers with self-reported risk for arboviral infection through the UNC Arboviral Traveler Study (IRB#08-0895). Plasma was isolated from whole blood by centrifugation and analyzed by virus-capture ELISA assay for binding to ZIKV and DENV 1-4 viruses. If antibody binding to any virus is observed, the neutralization titer was determined via FRNT_50_ neutralization. Plasma with 4-fold higher neutralization titers to one DENV serotype or ZIKV than all other titers was characterized as a primary infection (11). Previously characterized flaviviral positive serum samples were used as controls for ELISA and neutralization experiments.

### Viruses and Cells

ZIKV strain H/PF/2013 was obtained from the U.S. Centers for Disease Control and Prevention and used in all assays (25). DENV WHO reference strains DENV1 West Pac 74, DENV2 S16803, DENV3 CH54389, and DENV4 TVP-376 were initially obtained from Robert Putnak (Walter Reed Army Institute of Research, Silver Spring, Maryland, USA). A9E escape mutant viruses were isolated as previously described (11). For cell culture-based experiments and maintaining virus stocks, Vero (*Cercopithecus aethiops*) cells (ATCC CCL-81) were used. Vero cells were grown at 37°C and 5% CO_2_ in DMEM media supplemented with 5% fetal bovine plasma and L-glutamine. Virus stocks were titrated on Vero cells by plaque assay or focus-forming assay. All studies were conducted under biosafety level 2 containment.

### Memory B cell immortalization and sorting

The B11F monoclonal antibody was generated from donor DT 172 using the 6XL method of memory B cell immortalization (26). Briefly, PBMCs from donor DT 172 underwent CD22+ magnetic purification followed by flow cytometric sorting for CD19+CD27+IgM-class switched MBCs. These sorted MBCs were then transduced with the 6XL retrovirus and activated by incubation with CD40L expressing cells as well as human interleukin 21 to support antibody secretion and B cell proliferation (27). 6XL transduced MBCs then underwent flow cytometric sorting by green fluorescence protein (GFP) expression as a marker of transduction, into polyclonal cultures at 50 GFP+ cells per well on a 96-well plate using a BD FACSAria III.

### Memory B cell screening and monoclonal antibody generation

Polyclonal MBC cultures were screened by ELISA for binding to both DENV 1-4 and ZIKV. Cultures that were ZIKV positive were single-cell sorted on a BD FACSAria III, grown on CD40L and IL-21, and then screened as above after 4 weeks. ZIKV antibody positive monoclonal cultures were further screened by FRNT_50_ in Vero cells at 1:2 and 1:8 dilutions. Positive monoclonal cultures underwent RNA isolation and nested PCR for human IGH and IGL genes, followed by sequencing using previously described primers (28). Sequences were analyzed by IgBLAST (https://www.ncbi.nlm.nih.gov/igblast/) and compared to germline to determine VH and VL gene usage, V-(D)-J gene usage, CDR3 sequence, rate of somatic hypermutation and IgG isotype. The complete heavy chain and light chain V region sequences were then cloned into IgG1 (Genbank, FJ475055) and Igλ expression vectors (Genbank, FJ517647), respectively

Heavy and light chain vectors were verified by sequencing and transformed into DH5alpha cells (NEB). The transformed cells were grown and the plasmid was purified by midiprep (Macherey-Nagel). The purified plasmid DNA (both heavy and light chain) was transfected into a 30mL culture of HEK Expi293 cells (Thermo Scientific Expi293). Harvesting the culture after 5 days, the supernatant was affinity-purified with pre-equilibrated monoclonal antibody SelectSure resin in a gravity column. The column was washed with 1X PBS and eluted with 300mM sodium citrate pH=3.0, into six 475μL fraction tubes containing 25μL of 1M Tris pH=8.0.

### Capture ELISA

Monoclonal antibody binding to ZIKV, DENV, and subunit envelope antigens was determined by capture ELISA. A 96-well plate was coated with murine monoclonal antibody 4G2 (UNC Center for Structural Biology) for ZIKV and DENV antigens, anti-MBP monoclonal antibody (ProteinTech) for Zika EDI and EDIII antigens (29), or anti-His (Invitrogen) for recombinant ZVrecE80 antigen in 0.1M carbonate buffer pH 9.6. Plates were coated for 1 hour at 37°C, then washed with 1X TBS + 0.2% tween buffer using a plate washer (BioTek). 3% non-fat milk (in 1XTBS + 0.05% tween buffer) was used to block the plate. Antigens were added as follows: ZIKV diluted 1:1, EDI (200ng per well,), EDIII (200ng per well), ZVrecE80 (500ng per well) diluted in blocking buffer, incubated for 1 hour at 37°C then washed as done previously. B11F and control monoclonal antibodies were added to the plate at 100ng per well. We used an alkaline phosphatase conjugated goat anti-human IgG (Sigma) diluted 1:2500 in blocking buffer as a secondary antibody. Each incubation step was done for 1 hour at 37°C. PNPP substrate (Sigma) was added to develop the plate and absorbance was measured at 405nm using a plate reader (BioTek). All ELISA experiments were done in duplicate and as at least three independent experiments.

### Neutralization assay

Neutralization titers were determined by 96-well microFRNT as described previously (11). Briefly, Serial dilutions (1:4) of monoclonal antibody were mixed with 50-100 focus-forming units of virus 2% FBS DMEM media. The virus-antibody mixtures were incubated for 1 hour at 37°C and then transferred to a monolayer of Vero cells for infection for 40 hours with ZIKV (H/PF/2013). Cells were then fixed and permeabilized. Infected cells were stained with primary antibodies 4G2 (ATCC, HB-114) and 2H2 (UNC Center for Structural Biology) for a 1 hour at 37°C, washed and then incubated with horseradish peroxidase-conjugated goat anti-mouse secondary antibody (KPL) for 1 hour at 37°C. Foci were visualized with 50 µL of True Blue (KPL) and counted using a CTL ELISPOT reader. Cell only controls and ZIKV virus positive cell controls were also added to each plate. Neutralization experiments were done in duplicate and as at least three independent experiments.

### BOB Assay

BOB was performed as previously described (11). Briefly, a 96-well plate was coated with 4G2 at 100ng/well, and the plate was blocked with 3% non-fat milk diluted in 1X TBS + 0.05% tween. ZIKV was diluted 1:1 in blocking buffer and added to the plate. Monoclonal antibodies were serially diluted 1:4 in blocking buffer and added to the plate starting at 100ng/well. B11F was conjugated with alkaline phosphatase (Abcam) and added to the plate at 100ng/well. PNPP substrate (Sigma) was added and the 405nm absorbance was measured (BioTek).

### Escape Mutant Selection and Sequencing

ZIKV (MOI = 0.01) was incubated with different multiplications of the FRNT_50_ of B11F for 1 hour at 37°C. The virus-monoclonal antibody mixture was added (2 mL) to Vero cells in a 6-well plate (Greiner). After infecting the cells for 1 hour at 37°C, the supernatant was discarded and 1mL 2% FCS media (Gibco) + 1mL B11F diluted in 2% FCS was added to the plate. Wild-type ZIKV was passaged as a control in media alone alongside virus undergoing B11F selection, along with a cell only control. 150μL aliquots were taken at a 3-hour baseline, and at 24, 48, and 72 hours after infection for quantitative RT-PCR, and cytopathic effects were observed under a microscope for each timepoint. Three days after infection, 1 mL of the supernatant was passaged to a new plate of Vero cells + 1 mL 2% FCS media. RNA was isolated from the cell culture supernatants and converted to cDNA (NEB). The E gene of stock virus, passaged virus control and passaged virus + B11F were sequenced via RT-PCR. The PCR product was run on a 2% agarose gel, gel extracted and purified (Zymogen). The purified DNA product was submitted for sequencing. The ZIKV stock, passaged control and passaged + B11F were aligned via SnapGene. Mutations were observed and presented using PyMOL.

### Alanine Scanning Mutagenesis

Alanine scanning mutagenesis was carried out by Integral Molecular on an expression construct for ZIKV prM/E (strain ZikaSPH2015; UniProt accession # Q05320). Residues were mutagenized to create a library of clones, each with an individual point mutant (13). Residues were changed to alanine (with alanine residues changed to serine). The resulting ZIKV prM/E alanine-scan library covered 100% of target residues (672 of 672). Each mutation was confirmed by DNA sequencing, and clones were arrayed into 384-well plates, one mutant per well.

Cells expressing ZIKV E mutants were immunostained with the indicated antibodies and mean cellular fluorescence was detected using an Intellicyt flow cytometer. Mutations within critical clones were identified as critical to the monoclonal antibody epitope if they did not support reactivity of the mAb, but did support reactivity of other conformation-dependent monoclonal antibodies. This counter-screen strategy facilitates the exclusion of Env mutants that are globally or locally misfolded or that have an expression defect (30). Validated critical residues represent amino acids whose side chains make the highest energetic contributions to the monoclonal antibody-epitope interaction (31, 32).

## Acknowledgments

We would like to acknowledge and thank the donor volunteers of the UNC Arboviral Traveler Study. We would also like to thank Dr. Derek Carbaugh for help with analyzing sequences, and Dr. Ellen Young for sharing mAbs. This study was supported by NIH grants R01AI107731 (to AMDS), T32AI007151 (to AJM), T32AI055402 (to HAT), P20GM125498 and U01AI1141997 (to SAD), and CDC BAA 2017-N-18041 (to AMDS). AJM was also supported by the UNC School of Medicine Physician Scientist Training Award for this work. Some of the protein work done at UNC was supported by the National Cancer Institute of the National Institutes of Health under award number P30CA016086 (UNC Center for Structural Biology). The content is solely the responsibility of the authors and does not necessarily represent the official views of the National Institutes of Health. Cell sorting was done by the Flow Cytometry and Cell Sorting Facility at the Larner College of Medicine University of Vermont, (with thanks to Roxana del Rio-Guerra) and support by NIH grant S10ODO18175 (to Jonathan Boyson) and NIH grant P30GM118228. Sequencing work at UVM was done by the Vermont Integrated Genomics Resource and supported by NIH grant P30GM118228. NIH contract HHSN 75N93019C00073 (to BJD) supported alanine scanning studies.

## Author Contributions

SG performed experiments and contributed to writing the manuscript. HAT, BDM, NRG, SAD performed B cell sorting, screening and monoclonal antibody isolation. HAT and SAD also contributed to editing the manuscript and funding the project. AG, BJD and ED performed alanine scanning studies. AMD contributed to project funding and manuscript editing. AJM performed experiments and contributed to writing the manuscript.

## Competing Interests statement

AG, BJD and ED are employees of Integral Molecular.

## References

1. Zorrilla CD, Garcia Garcia I, Garcia Fragoso L, De La Vega A. 2017. Zika Virus Infection in Pregnancy: Maternal, Fetal, and Neonatal Considerations. J Infect Dis 216:S891–S896.

2. Lazear HM, Diamond MS. 2016. Zika Virus: New Clinical Syndromes and Its Emergence in the Western Hemisphere. J Virol 90:4864–4875.

3. Sridhar S, Luedtke A, Langevin E, Zhu M, Bonaparte M, Machabert T, Savarino S, Zambrano B, Moureau A, Khromava A, Moodie Z, Westling T, Mascarenas C, Frago C, Cortes M, Chansinghakul D, Noriega F, Bouckenooghe A, Chen J, Ng SP, Gilbert PB, Gurunathan S, DiazGranados CA. 2018. Effect of Dengue Serostatus on Dengue Vaccine Safety and Efficacy. N Engl J Med 379:327–340.

4. Henein S, Swanstrom J, Byers AM, Moser JM, Shaik SF, Bonaparte M, Jackson N, Guy B, Baric R, de Silva AM. 2017. Dissecting Antibodies Induced by a Chimeric Yellow Fever-Dengue, Live-Attenuated, Tetravalent Dengue Vaccine (CYD-TDV) in Naive and Dengue-Exposed Individuals. J Infect Dis 215:351–358.

5. Halstead SB. 2003. Neutralization and antibody-dependent enhancement of dengue viruses. Adv Virus Res 60:421–67.

6. Andrade DV, Harris E. 2018. Recent advances in understanding the adaptive immune response to Zika virus and the effect of previous flavivirus exposure. Virus Res 254:27–33.

7. Gallichotte EN, Young EF, Baric TJ, Yount BL, Metz SW, Begley MC, de Silva AM, Baric RS. 2019. Role of Zika Virus Envelope Protein Domain III as a Target of Human Neutralizing Antibodies. mBio 10.

8. Yu L, Wang R, Gao F, Li M, Liu J, Wang J, Hong W, Zhao L, Wen Y, Yin C, Wang H, Zhang Q, Li Y, Zhou P, Zhang R, Liu Y, Tang X, Guan Y, Qin CF, Chen L, Shi X, Jin X, Cheng G, Zhang F, Zhang L. 2017. Delineating antibody recognition against Zika virus during natural infection. JCI Insight 2.

9. Metz SW, Gallichotte EN, Brackbill A, Premkumar L, Miley MJ, Baric R, de Silva AM. 2017. In Vitro Assembly and Stabilization of Dengue and Zika Virus Envelope Protein Homo-Dimers. Sci Rep 7:4524.

10. Priyamvada L, Quicke KM, Hudson WH, Onlamoon N, Sewatanon J, Edupuganti S, Pattanapanyasat K, Chokephaibulkit K, Mulligan MJ, Wilson PC, Ahmed R, Suthar MS, Wrammert J. 2016. Human antibody responses after dengue virus infection are highly cross-reactive to Zika virus. Proc Natl Acad Sci U S A 113:7852–7.

11. Collins MH, Tu HA, Gimblet-Ochieng C, Liou GA, Jadi RS, Metz SW, Thomas A, McElvany BD, Davidson E, Doranz BJ, Reyes Y, Bowman NM, Becker-Dreps S, Bucardo F, Lazear HM, Diehl SA, de Silva AM. 2019. Human antibody response to Zika targets type-specific quaternary structure epitopes. JCI Insight 4.

12. Long F, Doyle M, Fernandez E, Miller AS, Klose T, Sevvana M, Bryan A, Davidson E, Doranz BJ, Kuhn RJ, Diamond MS, Crowe JE, Jr., Rossmann MG. 2019. Structural basis of a potent human monoclonal antibody against Zika virus targeting a quaternary epitope. Proc Natl Acad Sci U S A 116:1591–1596.

13. Sapparapu G, Fernandez E, Kose N, Bin C, Fox JM, Bombardi RG, Zhao H, Nelson CA, Bryan AL, Barnes T, Davidson E, Mysorekar IU, Fremont DH, Doranz BJ, Diamond MS, Crowe JE. 2016. Neutralizing human antibodies prevent Zika virus replication and fetal disease in mice. Nature 540:443–447.

14. Niu X, Zhao L, Qu L, Yao Z, Zhang F, Yan Q, Zhang S, Liang R, Chen P, Luo J, Xu W, Lv H, Liu X, Lei H, Yi C, Li P, Wang Q, Wang Y, Yu L, Zhang X, Bryan LA, Davidson E, Doranz JB, Feng L, Pan W, Zhang F, Chen L. 2019. Convalescent patient-derived monoclonal antibodies targeting different epitopes of E protein confer protection against Zika virus in a neonatal mouse model. Emerg Microbes Infect 8:749–759.

15. Ravichandran S, Hahn M, Belaunzaran-Zamudio PF, Ramos-Castaneda J, Najera-Cancino G, Caballero-Sosa S, Navarro-Fuentes KR, Ruiz-Palacios G, Golding H, Beigel JH, Khurana S. 2019. Differential human antibody repertoires following Zika infection and the implications for serodiagnostics and disease outcome. Nat Commun 10:1943.

16. Stettler K, Beltramello M, Espinosa DA, Graham V, Cassotta A, Bianchi S, Vanzetta F, Minola A, Jaconi S, Mele F, Foglierini M, Pedotti M, Simonelli L, Dowall S, Atkinson B, Percivalle E, Simmons CP, Varani L, Blum J, Baldanti F, Cameroni E, Hewson R, Harris E, Lanzavecchia A, Sallusto F, Corti D. 2016. Specificity, cross-reactivity, and function of antibodies elicited by Zika virus infection. Science 353:823–6.

17. Wang J, Bardelli M, Espinosa DA, Pedotti M, Ng TS, Bianchi S, Simonelli L, Lim EXY, Foglierini M, Zatta F, Jaconi S, Beltramello M, Cameroni E, Fibriansah G, Shi J, Barca T, Pagani I, Rubio A, Broccoli V, Vicenzi E, Graham V, Pullan S, Dowall S, Hewson R, Jurt S, Zerbe O, Stettler K, Lanzavecchia A, Sallusto F, Cavalli A, Harris E, Lok SM, Varani L, Corti D. 2017. A Human Bi-specific Antibody against Zika Virus with High Therapeutic Potential. Cell 171:229–241 e15.

18. Metz SW, Thomas A, Brackbill A, Forsberg J, Miley MJ, Lopez CA, Lazear HM, Tian S, de Silva AM. 2019. Oligomeric state of the ZIKV E protein defines protective immune responses. Nat Commun 10:4606.

19. Barba-Spaeth G, Dejnirattisai W, Rouvinski A, Vaney MC, Medits I, Sharma A, Simon-Loriere E, Sakuntabhai A, Cao-Lormeau VM, Haouz A, England P, Stiasny K, Mongkolsapaya J, Heinz FX, Screaton GR, Rey FA. 2016. Structural basis of potent Zika-dengue virus antibody cross-neutralization. Nature 536:48–53.

20. Hasan SS, Miller A, Sapparapu G, Fernandez E, Klose T, Long F, Fokine A, Porta JC, Jiang W, Diamond MS, Crowe JE, Jr., Kuhn RJ, Rossmann MG. 2017. A human antibody against Zika virus crosslinks the E protein to prevent infection. Nat Commun 8:14722.

21. Robbiani DF, Bozzacco L, Keeffe JR, Khouri R, Olsen PC, Gazumyan A, Schaefer-Babajew D, Avila-Rios S, Nogueira L, Patel R, Azzopardi SA, Uhl LFK, Saeed M, Sevilla-Reyes EE, Agudelo M, Yao KH, Golijanin J, Gristick HB, Lee YE, Hurley A, Caskey M, Pai J, Oliveira T, Wunder EA, Jr., Sacramento G, Nery N, Jr., Orge C, Costa F, Reis MG, Thomas NM, Eisenreich T, Weinberger DM, de Almeida Arp, West AP, Jr., Rice CM, Bjorkman PJ, Reyes-Teran G, Ko AI, MacDonald MR, Nussenzweig MC. 2017. Recurrent Potent Human Neutralizing Antibodies to Zika Virus in Brazil and Mexico. Cell 169:597–609 e11.

22. Wang Q, Yang H, Liu X, Dai L, Ma T, Qi J, Wong G, Peng R, Liu S, Li J, Li S, Song J, Liu J, He J, Yuan H, Xiong Y, Liao Y, Li J, Yang J, Tong Z, Griffin BD, Bi Y, Liang M, Xu X, Qin C, Cheng G, Zhang X, Wang P, Qiu X, Kobinger G, Shi Y, Yan J, Gao GF. 2016. Molecular determinants of human neutralizing antibodies isolated from a patient infected with Zika virus. Sci Transl Med 8:369ra179.

23. Zhang S, Kostyuchenko VA, Ng TS, Lim XN, Ooi JSG, Lambert S, Tan TY, Widman DG, Shi J, Baric RS, Lok SM. 2016. Neutralization mechanism of a highly potent antibody against Zika virus. Nat Commun 7:13679.

24. Vita R, Mahajan S, Overton JA, Dhanda SK, Martini S, Cantrell JR, Wheeler DK, Sette A, Peters B. 2019. The Immune Epitope Database (IEDB): 2018 update. Nucleic Acids Res 47:D339–D343.

25. Baronti C, Piorkowski G, Charrel RN, Boubis L, Leparc-Goffart I, de Lamballerie X. 2014. Complete coding sequence of zika virus from a French polynesia outbreak in 2013. Genome Announc 2.

26. Kwakkenbos MJ, Diehl SA, Yasuda E, Bakker AQ, van Geelen CM, Lukens MV, van Bleek GM, Widjojoatmodjo MN, Bogers WM, Mei H, Radbruch A, Scheeren FA, Spits H, Beaumont T. 2010. Generation of stable monoclonal antibody-producing B cell receptor-positive human memory B cells by genetic programming. Nat Med 16:123–8.

27. Diehl SA, Schmidlin H, Nagasawa M, van Haren SD, Kwakkenbos MJ, Yasuda E, Beaumont T, Scheeren FA, Spits H. 2008. STAT3-mediated up-regulation of BLIMP1 Is coordinated with BCL6 down-regulation to control human plasma cell differentiation. J Immunol 180:4805–15.

28. Ho IY, Bunker JJ, Erickson SA, Neu KE, Huang M, Cortese M, Pulendran B, Wilson PC. 2016. Refined protocol for generating monoclonal antibodies from single human and murine B cells. J Immunol Methods 438:67–70.

29. Premkumar L, Collins M, Graham S, Liou GA, Lopez CA, Jadi R, Balmaseda A, Brackbill JA, Dietze R, Camacho E, De Silva AD, Giuberti C, Dos Reis HL, Singh T, Heimsath H, Weiskopf D, Sette A, Osorio JE, Permar SR, Miley MJ, Lazear HM, Harris E, de Silva AM. 2018. Development of Envelope Protein Antigens To Serologically Differentiate Zika Virus Infection from Dengue Virus Infection. J Clin Microbiol 56.

30. Davidson E, Doranz BJ. 2014. A high-throughput shotgun mutagenesis approach to mapping B-cell antibody epitopes. Immunology 143:13–20.

31. Bogan AA, Thorn KS. 1998. Anatomy of hot spots in protein interfaces. J Mol Biol 280:1–9.

32. Lo Conte L, Chothia C, Janin J. 1999. The atomic structure of protein-protein recognition sites. J Mol Biol 285:2177–98.

